# Nitrate Promoting Hair Growth Through Enhancing Wnt/β-catenin and Downregulating TGF-β/Smad/BMP Signaling Pathways in C57BL/6 Mice

**DOI:** 10.1101/2024.11.02.621646

**Authors:** Yanjie Lan, Shenglan Li, Jiachen Wang, Ling Duan, Mengqian Huang, Xin Yang, Feng Chen, Chunmei Zhang, Songling Wang, Wenbin Li

## Abstract

Nitrate found widely distributed in vegetables and fruits. It has been proved to have many physiological functions through nitrate-nitrite-NO pathway, such as attenuate oxidative stress, enhance the skeletal muscle and tone skin vascular. But the function of nitrate in regulating hair growth is unclear. Here, we evaluated potential roles of nitrate on hair growth and its mechanisms by *in vivo* models. Mice treated with 10mM nitrate supplementation significantly promoting hair growth, along with an increase in the number of hair follicles and thickness of dermis. We further demonstrated that Wnt3a and β-catenin were highly expressed in the nitrate-treated groups, while the expression of TGF-β1 was attenuated. Moreover, nitrate also increased the expression levels of growth factors (Fgf, Igf-1, and Vegf). Our data indicated that nitrate supplementation could effectively accelerate hair regrowth through activating the Wnt/β-catenin and inhibiting TGF-β1/Smad/BMP signaling pathway. Therefore, these findings provide support for a potential therapeutic role of nitrate in preventing and treating alopecia.

## Introduction

Hair loss has become a common worldwide problem currently. It has been found that hair loss progresses owing to multiple factors such as stress, hormonal imbalance, diet, and intensification of pollution, along with several genetic factors[1]. Hair grows from hair follicles, which undergo a continuous cyclical pattern characterized by phases of regeneration (anagen), regression (catagen), rest (telogen), and shedding of old hair fiber (exogen)[2, 3]. Normally, hair follicles are contracted after the anagen phase and the hair shaft falls out during the catagen or telogen phase. As the follicles begin a new anagen phase, they grow back to their original size and produce new hair of a normal thickness[4, 5]. If the cycle is not properly regulated, resulting in hair loss[4, 6]. At present, only minoxidil and finasteride these two effective drugs approved by the US Food and Drug Administration (FDA) for promoting hair growth and preventing hair loss[7]. Given that the application of both minoxidil and finasteride is limited due to the side effects caused by long-term administration, such as sexual dysfunction, facial hairiness and scalp stimulation[8, 9]. Thus, it is crucial to seek an effective therapeutic alternatives or products for promote hair growth.

Nitrate, serves as a natural constituent of the human diet. Fruits and vegetables, including beetroot juice, spinach, and celery, are major contributors to nitrate[10, 11]. It is clear that nitrate can be continuously metabolized to NO and other bioactive nitrogen oxides through nitrate-nitrite-NO pathway, which boost various biological activities of nitrate such as attenuate oxidative stress, improve endothelial function, and enhance the skeletal muscle contractile properties[12, 13]. Studies found that NO exhibits positive functions and plays an essential role in the physiology of the hair growth. Wolf R., *et al*., suggested NO might act as a signaling molecule in human dermal papilla cells and involved in the regulation of hair follicle activity[14]. In addition, the topical application of non-thermal plasma in the dermis of mouse dorsal skin was found to generate large amounts of exogenous NO to improved hair growth[15]. More importantly, researchers suggested dietary nitrate supplementation can increase substrates for oxygen-independent NO synthesis in the skin interstitial fluid[16]. Accordingly, Dietary nitrate as one effective way of delivering nitrite and NO systemically, the function of it in regulating hair growth have not been elucidated yet.

A variety of signaling pathways have been implicated in regulating the growth, development and cycle of hair follicles in humans and mice, including Wnt/β-catenin, transforming growth factor-β/bone morphogenic protein (TGF-β/BMP) and Sonic hedgehog (Shh)[17-19] and so on. Targeting these multicomponent signaling of hair growth regulation would be a rational approach for developing novel therapeutics for the treatment of hair loss. Chemical components[20-22], plant extracts[23-26] and intestine microorganism[27] are reported to regulate some of these signaling pathways in promoting hair growth *in vivo* and *in vitro*. It is worth noting that some studies proposed that phytochemicals are more effective than conventional hair loss regimens such as minoxidil and finasteride[28, 29]. Moreover, research on hair growth therapies has mostly focused on direct topical treatments, whereas oral treatments have remained largely unexplored[30].

In present study, we investigated the function of dietary nitrate in hair growth as well as its potential mechanism of action by using *in vivo* models. The hair growth was evaluated after administering dietary nitrate for four weeks. Gene and protein expression of hair growth-related factors in the skin tissues of nitrate-treated C57BL/6 mice were evaluated. Our study revealed that nitrate showed a hair growth promoting activity via modulation of Wnt/β-catenin and TGF-β signaling pathways *in vivo*. We propose that dietary nitrate may be a promising alternative therapy for hair loss.

## Materials and methods

### Animals

Six-week-old female C57BL6 mice (Spf animals, Beijing, China) were used for the present study. All animals were maintained at 23 ± 2 °C, a relative humidity of 50 ± 5%, a 12:12 h light-dark cycle, and with free access to water and pellet food. This study was approved by the animal ethics committee of Capital Medical University (Beijing, china, approval no. 202202004).

### Chemicals and study design

Nitrate used in the experiment were purchased from Sigma (USA). Nitrate were diluted to 1 M as the stock dilution and 10 mM as the usage dilution. After a one-week of acclimation, mice were anesthetized using 1.25% 2,2,2-tribromoethanol by intraperitoneal injection, and then remove the hair on the back of all mice. All mice were randomized into two groups: control group (n=5) and nitrate experimental group (n=5). 10 mM nitrate was administered orally in the drinking water to mice in nitrate group for 4 weeks, while mice in control group were given regular drinking water. The changes in hair growth were recorded on days 0, 7, 14 and 21. Skin samples resected from the back of mice in two group were collected approximately 2×4 cm for analyses. All animal experiments were carried out according to the NIH Guideline for the Care and Use of Laboratory Animals.

### Assessing the degree of hair growth

The degree of hair growth of mice in different groups was evaluated according to the criteria listed in Table 1 [31] .

### Hematoxylin and eosin (H&E) stain

Skin samples collected on days 0, 7, 14 and 21 were fixed in 4% PFA for 24 h, dehydrated in graded ethanol (50%, 75%, 85%, 95%, 100%), hyalinized by dimethylbenzene, filled with paraffin and sectioned into 4 μm. Then tissues were stained with H&E according to the manufacturer’s instructions.

### Immunohistochemistry assays

Formalin-fixed and paraffin-embedded skin tissue sections were deparaffinized, rehydrated, and heated in sodium citrate buffer (pH 6.0) or EDTA Antigen Retrieval Solution (pH 8.0) prior to retrieval antigen. Then blocking endogenous peroxidase for 30min, and blocked with 5% normal goat serum for 60min at room temperature. Afterwards, tissue sections were incubated with primary antibody overnight at 4°C. Followings are primary antibodies used here: anti-β-catenin (Proteintech, USA; 1:1000 dilution), anti-Ki67 (Abclonal, China; 1:200 dilution), anti-wnt3a (Abcam, USA, 1:200 dilution) and anti-TGF-β1(Proteintech, USA; 1:200 dilution). HRP-conjugated goat anti-rabbit antibody (Zhongshan Goldenbridge Biotechnology, Beijing, China) was used as the secondary antibody to incubate with the sections for 60min at room temperature. Finally, immunoreaction was visualized using the diaminobenzidine tetrahydrochloride-h2o2 method. The sections were mounted and observed under the inverted microscopy (Zeiss Vert.A1, German)

### Immunofluorescence assays

The dorsal skin sections were permeabilized with 0.3% Triton X-100 for 30 min at room temperature and blocked in 5% normal goat serum for 60 min. Then they were incubated with primary antibodies overnight at 4°C. Followings are primary antibodies used here: anti-β-catenin (Proteintech, USA; 1:1000 dilution) and anti-wnt3a (Abcam, USA, 1:200 dilution). Then sections were probed with secondary antibody: FITC-conjugated goat anti-rabbit IgG or Alexa Fluor® 594 conjugated goat anti-rabbit IgG (Zhongshan Goldenbridge Biotechnology, Beijing, China) for 60min at room temperature. The nuclei were stained with 4′,6-diamidino-2-phenylindole and counterstained with DAPI (Solarbio, Beijing, China). Sections were covered with mounting medium, and images of tissue sections were obtained using a fluorescence microscope (Invitrogen EVOS FL Auto2).

### Quantitative real-time PCR

Total RNA was extracted from dorsal skin tissues of mice using the Trizol-chloroform method. The total RNA concentration was determined using a by Thermo Scientific^TM^ NanoDrop^TM^ One (Thermo Scientific, USA). Further, total RNA was used as a template for cDNA synthesis using the cDNA synthesis kit (TransScript® All-in-One First-Strand cDNA Synthesis SuperMix for qPCR). Quantitative real-time PCR was performed using the PowerUP^TM^ SYBR^TM^ Green Supermix (Applied Biosysterm^TM^, USA), in which the expression of target genes was normalized to that of GAPDH. The primers used are presented in Table 2.

### Western blot

Skin samples were homogenized with RIPA lysis buffer containing phosphatase and protease inhibitor. Total protein concentration was quantified by BCA assay. 60 μg concentrations of protein were loaded on 10% SDS-polyacrylamide gel for electrophoresis. The proteins in the gel were transferred to polyvinylidene difluoride membrane. Then, the membrane was blocked with 5% skimmed milk for 1h at room temperature. After that, the membrane was incubated with primary antibodies at 4°C for 6-8h. After washing, the membrane was incubated with secondary antibodies for 1h at room temperature. Western blot detection was performed using a chemiluminescence reagent. The primary antibodies used were as follows: anti-β-catenin (Proteintech, USA; 1:1000 dilution), anti-Cyclin D1 (Cell Signaling Technology, USA; 1:1000 dilution), anti-TGF-β1 (Proteintech, USA; 1:1000 dilution) and anti-β-actin (CST, USA; 1:1000 dilution). HRP-conjugated anti-rabbit IgG (Zhongshan Goldenbridge Biotechnology, Beijing, China) was used as the secondary antibodies. β-actin was used as the reference primary antibody.

### Statistical analysis

All data were presented as mean ± standard deviation. P < 0.05 was considered statistically significant. Statistical analysis was performed by GraphPad Prism software (8.4.0). When parameters followed Gaussian distribution, Student’s t test was used for two groups’ analyses and one-way ANOVA was used for comparing more than two groups to evaluate the statistical significance. The significance of differences was determined based on P values, where *P < 0.05; **P < 0.01; ***P < 0.001; ****P < 0.0001. Sample size was determined according to experience or the previously published papers.

## Results

### Nitrate promoted hair growth in C57BL/6 mice

The mouse model of depilation-induced hair regeneration was constructed to detect the effects of nitrate on hair growth. Then, mice were randomly divided into two groups and treated with oral administration of nitrate (10 mM) or vehicle, respectively. The oral administration of nitrate did not result in any observable skin disease, toxicity and measurements of body weight in mice showed no abnormal changes (Figure S1A-B). Photographs were taken to visually confirm the hair growth pattern of the mouse dorsal skin (Figure 1A). Both groups of mice showed pink skin after depilation on the first day, which confirmed that most of their hair was in the resting phase. On the 7th day, the dorsal skin began to appear gray in the group of mice with oral administration of nitrate, which indicated that nitrate-treated mice entered into the anagen growth phase of the hair cycle faster than control mice. After 10 days of treatment, the nitrate group was in a darker shade of gray, while the control group was pale gray. After 14 days, hair growth was significantly improved in the nitrate group, whereas there could still observe the bare patches on dorsal skin in controls. Consistently, the hair growth score was calculated and showed that nitrate group had higher score than controls after 7, 10 and 14 days of treatment (Figure 1B-C). At 21th days, mice in nitrate group exhibited full hair growth. Taken together, these results showed that nitrate supplementation could effectively promote hair growth.

**Figure 1.**
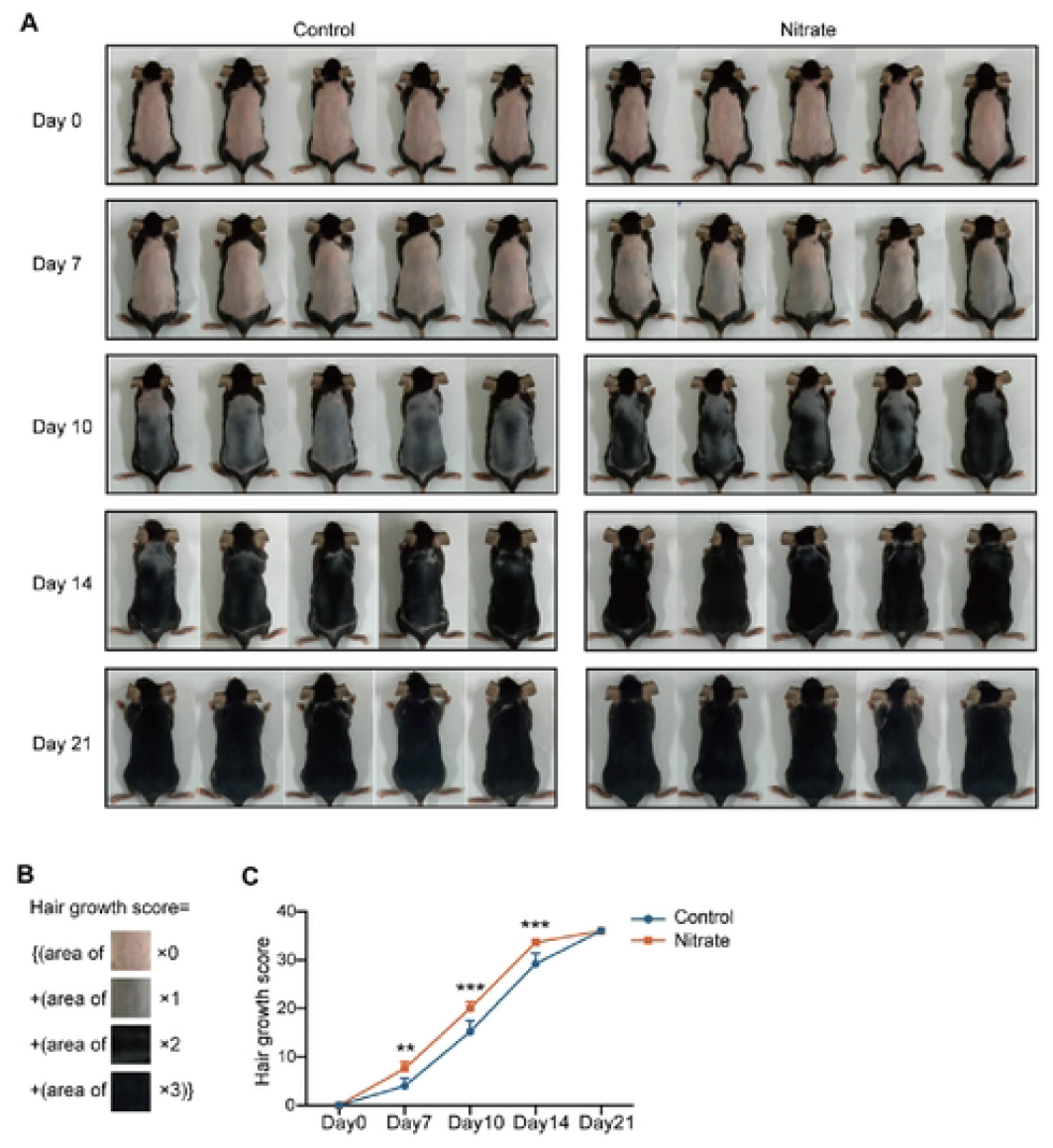
Hair growth-promoting effects of oral nitrate in C57BL/6 mice. (A) Photographs of C57BL/6 mice dorsal skin taken on days 0, 7, 10, 14, and 21 after depilated and repetitively treated with oral administration of nitrate (10mM) or vehicle. (B) Image of hair growth index calculated with scores. (C) Quantification of hair growth scores. All data are presented as the mean ± SD (n=4). * p < 0.05, ** p < 0.001, and *** p < 0.0001.

### Nitrate increased the number of hair follicles and thickness of dermis *in vivo*

Skin specimens of the depilation area of two groups of mice were collected after 0, 7, 14 and 21 days of treatment. H&E staining was performed to investigate the effect of nitrate on the development of hair follicles and skin thickness. On day 7, the hair bulbs of the oral nitrate-treated mice began to become plump and migrate downward, revealing that the hair follicles were transformed from the telogen phase to the anagen phase. At this phase, the number of hair follicles was increased in mice treated with nitrate compared with controls (Figure 2A-B). After 14 days, the hair bulbs were located at the dermal-subcutaneous junction, which indicating that the hair follicles were in the anagen phase. The number of hair follicles was in a significant increase in nitrate group, while the hair follicles of the controls were sparse (Figure 2A-B). Moreover, hair follicles formed more melanin in this phase in mice skin treated with oral nitrate. At the 21st day, mice in nitrate group had a smaller number of hair follicles than that in control group (Figure 2A-B). In addition, compared to the control, we also observed that mice treat with nitrate increased the thickness of dermis (Figure 2A and 2C). These observations suggested that the nitrate supplementation could improve hair growth by increasing the number of hair follicles and thickness of dermis, especially in the 7-14 days after treatment.

**Figure 2.**
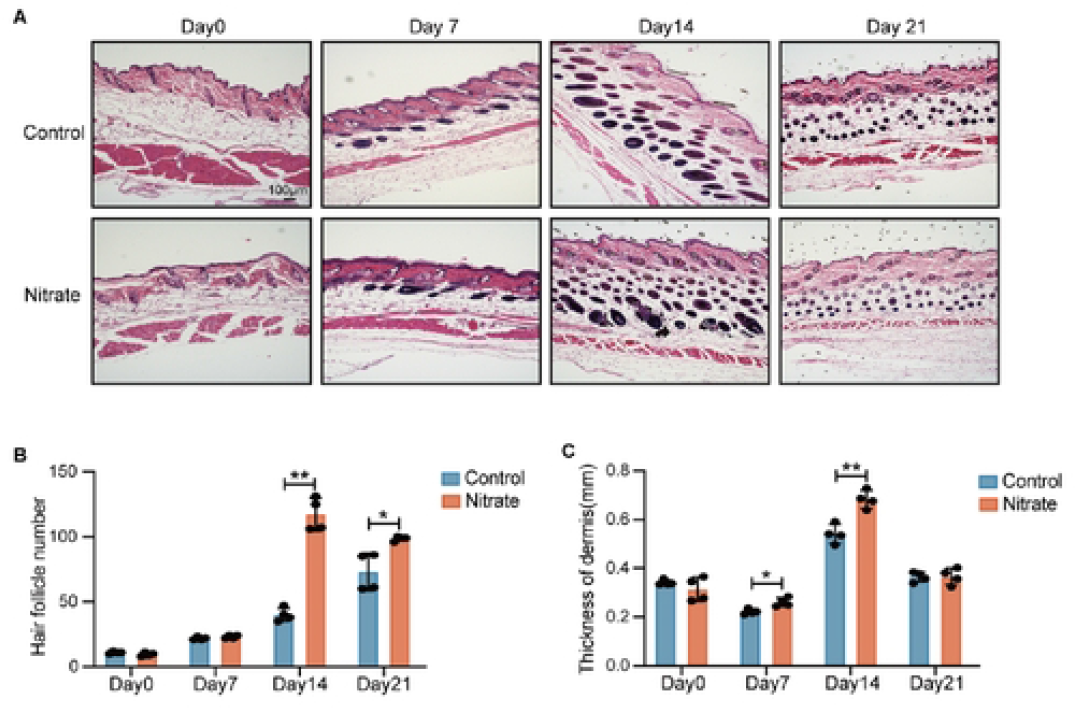
Nitrate accelerated hair growth and alters the number of hair follicles and thickness of dermis. (A) H&E staining images (40×) of longitudinal skin sections from mice treated with nitrate (10mM) or vehicle at days 0, 7, 14, and 21. (B) The number of hair follicles of the dorsal skin from mice treated with nitrate (10mM) or vehicle at days 0, 7, 14, and 21. (C) The **t**hickness of dermis was measured in mice treated with nitrate (10mM) or vehicle at days 0, 7, 14, and 21. All data are presented as the mean ± SD (n=4). * p < 0.05, ** p < 0.001, and *** p < 0.0001.

### Nitrate promoted hair growth by enhancing the Wnt/β-catenin signaling pathway

Among various signaling pathways, Wnt/β-catenin signaling is an essential pathway in regulating hair morphogenesis and hair growth. To clarify the effect of nitrate on the activation of Wnt/β-catenin signaling in vivo, specimens of the dorsal skin were collected on days 0, 7, 14 and 21 for analyses. Notably, immunohistochemistry (IHC) staining revealed an increase in β-catenin level of nitrate group compared to those observed in the control group after treatment of 7 and 14 days (Figure 3A). According to immunofluorescence assays (Figure 3B), nitrate significantly increased the fluorescence signals of β-catenin in hair follicles compared to controls on the 7th and 14th day. We also examined the expressions these markers by western blot analysis. β-catenin, and its downstream molecules such as CyclinD1 was trended upwards by nitrate treatments (Figure 3C). Consistently, RT-PCR results showed nitrate treatments markedly upregulated the mRNA expression of β-catenin, Lef-1 and Gsk-3β (Figure S2A-C). Similarly, Sonic hedgehog (Shh) signaling also mediates β-catenin activity to direct hair follicle morphogenesis and hair cycle initiation. Compared with control, nitrate treatments significantly upregulated the mRNA expression of Shh at day 7 and day 14 (Figure 3D). In addition, Ki67, as a marker of both the β-catenin and proliferation marker, was also increased in the nitrate-treated mice compared with controls by IHC staining at day 7 and day 14 (Figure 3E). The mRNA expression of Ki67 was also increased in nitrate-treated mice (Figure S2D).

**Figure 3.**
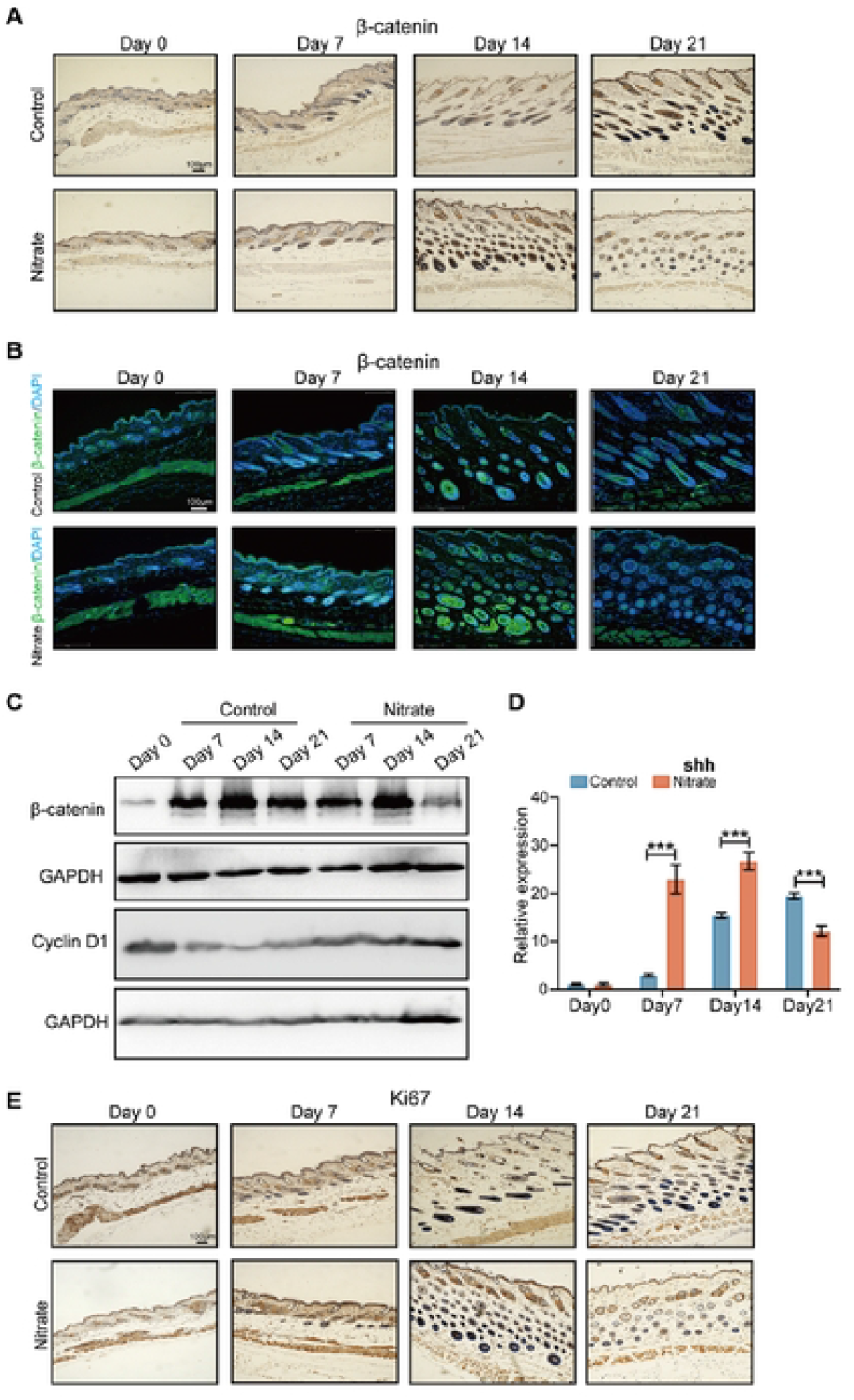
Nitrate activated β-catenin and its downstream molecules in mice in promoting hair growth. (A) Representative immunohistochemical images showing the expressions of β-catenin in the dorsal skin samples from mice treated with vehicle (control group) or nitrate (10mM) on days 0, 7, 14 and 21. Scale bar = 50 μm. (B) Immunofluorescence staining of β-catenin (green fluorescence) and DAPI (blue fluorescence) of hair follicles in skin was performed in mice treated with nitrate (10mM) or vehicle at days 0, 7, 14, and 21. Scale bars = 100 μm. (C) Protein expression level of β-catenin and cyclin D1 in the mouse skin tissue were detected by western blot; (D) Relative mRNA expression of Shh in mice treated with nitrate (10mM) or vehicle at days 0, 7, 14, and 21; (E) Immunohistochemistry analysis of the expressions of Ki67 in mice treated with nitrate (10mM) or vehicle at days 0, 7, 14, and 21. Scale bar: 50 μm. All data are presented as the mean ± SD (n=4). * p < 0.05, ** p < 0.001, and *** p < 0.0001.

To further assess whether Wnt signaling is involved in the hair growth-stimulating activity of nitrate, we detected the mRNA expression of critical genes related to Wnt signaling. The expression of wnt4a, wnt5a, wnt5b and wnt7a in the nitrate-treated groups was in a significant upregulation at day 7 and day 14 (figure S3A-D). Wnt3a was an activate factor of β-catenin and played a significant role in promoting hair growth by regulating the dermal papilla cells. At the protein level, nitrate also induced a marked increasing of Wnt3a as determined by immunohistochemistry and immunofluorescence analyses (Figure 4A-B). Similarly, the mRNA expression of Wnt3a was increased in the nitrate-treated groups (Figure 4C). Specifically, Wnt10b is prominently upregulated in the hair follicle epithelium at the onset of the anagen phase. We also found mice treated with nitrate exhibited higher mRNA expression levels at day 7 and 14. (Figure 4D). These results suggested that nitrate supplementation promotes the telogen-to-anagen transition by enhancing the Wnt/β-catenin signaling pathway

**Figure 4.**
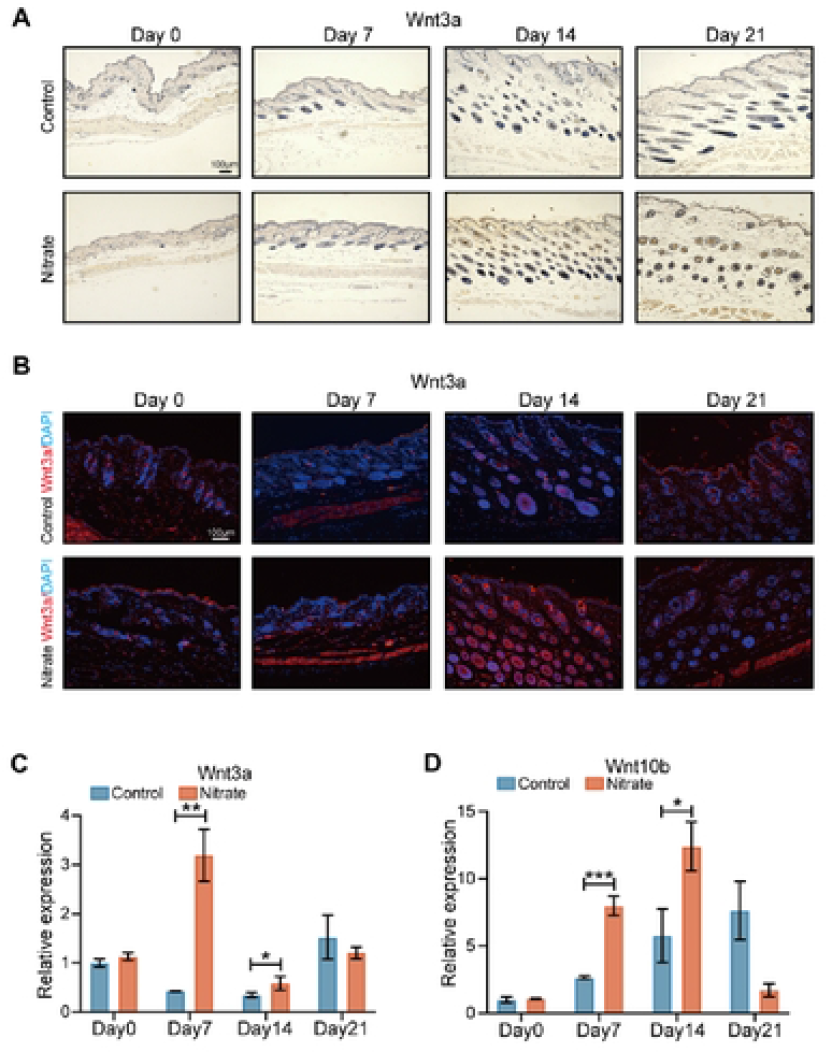
Nitrate upregulated wnt3a and critical genes related to Wnt signaling in mice in promoting hair growth. (A) Representative immunohistochemical images showing the expressions of wnt3a in the dorsal skin samples from mice treated with vehicle (control group) or nitrate (10mM) on days 0, 7, 14 and 21. Scale bar = 50 μm. (B) Immunofluorescence staining of wnt3a (red fluorescence) and DAPI (blue fluorescence) of hair follicles in skin was performed in mice treated with nitrate (10mM) or vehicle at days 0, 7, 14, and 21. Scale bars = 100 μm. (C-D) Relative mRNA expression of Wnt3a and Wnt10b in mice treated with nitrate (10mM) or vehicle at days 0, 7, 14, and 21; (C) Wnt3a, (D) Wnt10b. All data are presented as the mean ± SD (n=4). * p < 0.05, ** p < 0.001, and *** p < 0.0001.

### Nitrate deregulated TGF-β/Smad/BMP signaling in hair follicles

TGF-β1 is a negative regulator of hair growth, which can lead to the induction of catagen in hair follicles. To verify whether nitrate promoted hair growth through inhibiting TGF-β signaling axis, we detected the TGF-β1 levels by IHC. We found that the level of TGF-β1 was reduced after nitrate treatments of 7 and 14 days (Figure 5A). Simultaneously, we further confirmed its level in mRNA and protein level, and found the mRNA and protein expression of TGF-β1 in the dorsal skin of mice were reduced at day 7 and 14 (Figure 5B-C). Studies have shown that TGF-β/Smad signaling pathway was associated with cell cycle regulation in hair follicular cells. We further determined whether nitrate can inhibit the activation of the Smad2. The results showed that mice treatment with nitrate resulted in a significant decrease in Smad2 level as expected at day 7 and 14 (Figure 5D). Bone morphogenetic proteins (BMPs) belong to the TGF-β family and plays pivotal roles in controlling the initiation of the hair follicle growth phase by stopping cell proliferation and differentiation during epidermal development. Furthermore, we measured the mRNA expression of Bmp2, Bmp4, and Bmp6 by RT-PCR. Results showed that the mRNA levels of Bmp2 was decreased in mice treated with nitrate at day 7, and the mRNA levels of Bmp6 was decreased at day 14 in nitrate treated group. Whereas there was no significant difference in the mRNA levels of Bmp4 between two groups of mice (Figure 5E-G). Overall, these results suggested that nitrate supplementation might inhibit TGF-β/Smad/BMP signaling in modulating hair growth in mice.

**Figure 5.**
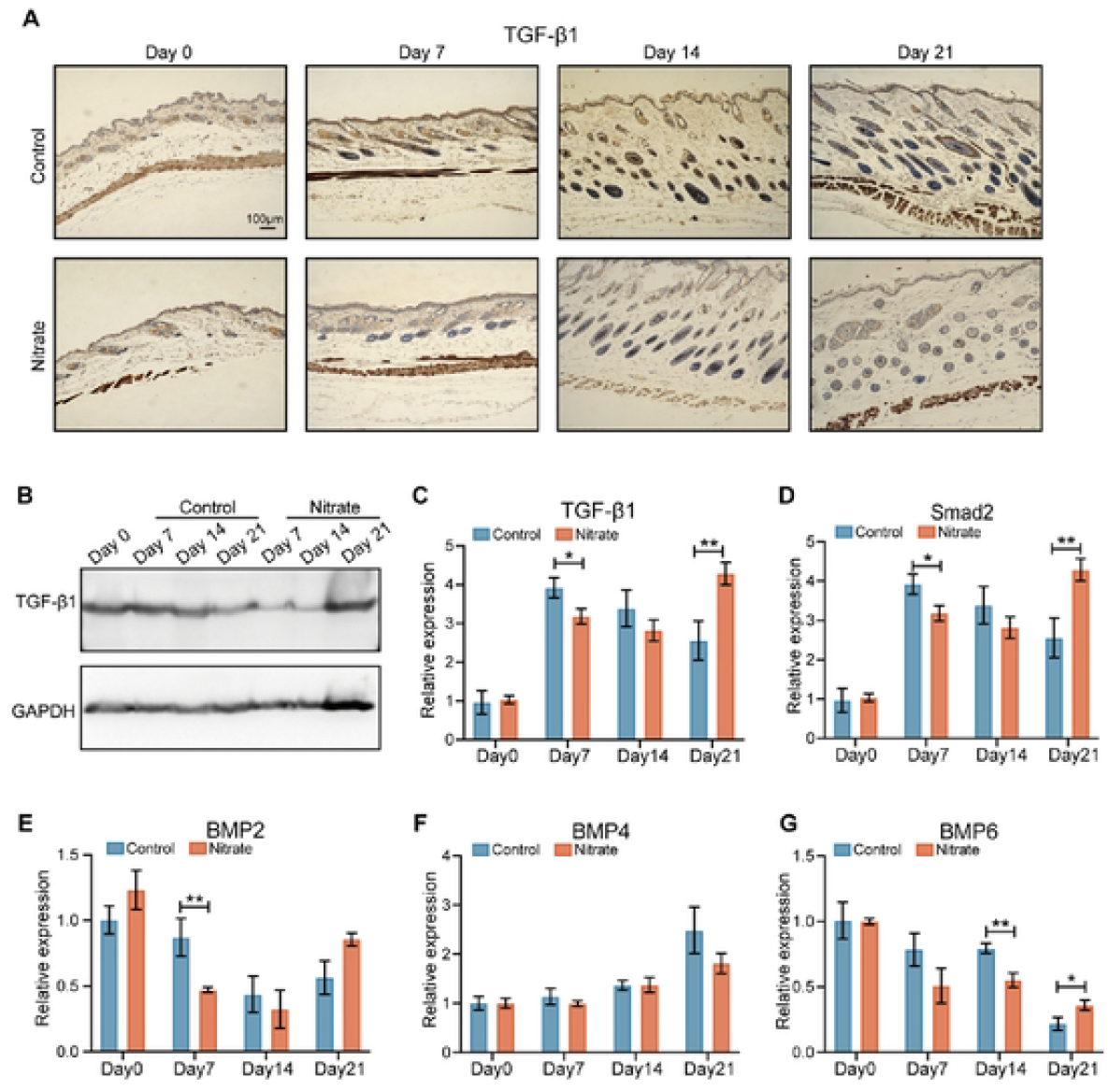
Nitrate suppressed the expression of TGF-β1 in mice to induce hair growth. (A) Representative immunohistochemical images showing the expressions of TGF-β1 in the dorsal skin samples from mice treated with vehicle (control group) or nitrate (10mM) on days 0, 7, 14 and 21. Scale bar = 50 μm. (B) Protein expression level of TGF-β1 in the mouse skin tissue were detected by western blot; (C) Relative mRNA expression of TGF-β1 in mice treated with nitrate (10mM) or vehicle at days 0, 7, 14, and 21; (D-E) Relative mRNA expression of BMP2, BMP4 and BMP6 in mice treated with nitrate (10mM) or vehicle at days 0, 7, 14, and 21; (D) BMP2, (E) BMP4, (F) BMP6. All data are presented as the mean ± SD (n=4). * p < 0.05, ** p < 0.001, and *** p < 0.0001.

### Nitrate increased the expression of Vegf, Fgf2 and Igf-1 in vivo

Vascular endothelial growth factor (VEGF), fibroblast growth factor (FGF) and insulin-like growth factor-1 (IGF-1) are believed to promote various stages of the hair cycle and have pivotal roles in hair growth. IGF-1 is an important growth factor that regulates hair growth. The mRNA expression of Igf-1 was increasing at day 7 and day 14 in nitrate group (Figure 6A). VEGF promotes angiogenesis around hair follicles and induces hair anagen. To investigate the involvement of VEGF signaling in hair growth by nitrate treatment, we analyzed the mRNA expression of Vegfa and Vegfb by RT-PCR. Results revealed that both of them were increased at day 7 and day 14 in nitrate group (Figure 6B-C). Moreover, FGF was an important growth factor known to stimulate hair growth. We also measured the mRNA expression of Fgf2, Fgf7 and Fgf10. Nitrate treatment upregulated the Fgf2 mRNA expression compared to that observed in controls, while there was no significant difference between two groups in the mRNA expression of Fgf7 and Fgf10 (Figure 6D-F). Therefore, our results suggested that nitrate supplementation might increase the expression of VEGF, FGF2 and IGF in promoting hair growth.

**Figure 6.**
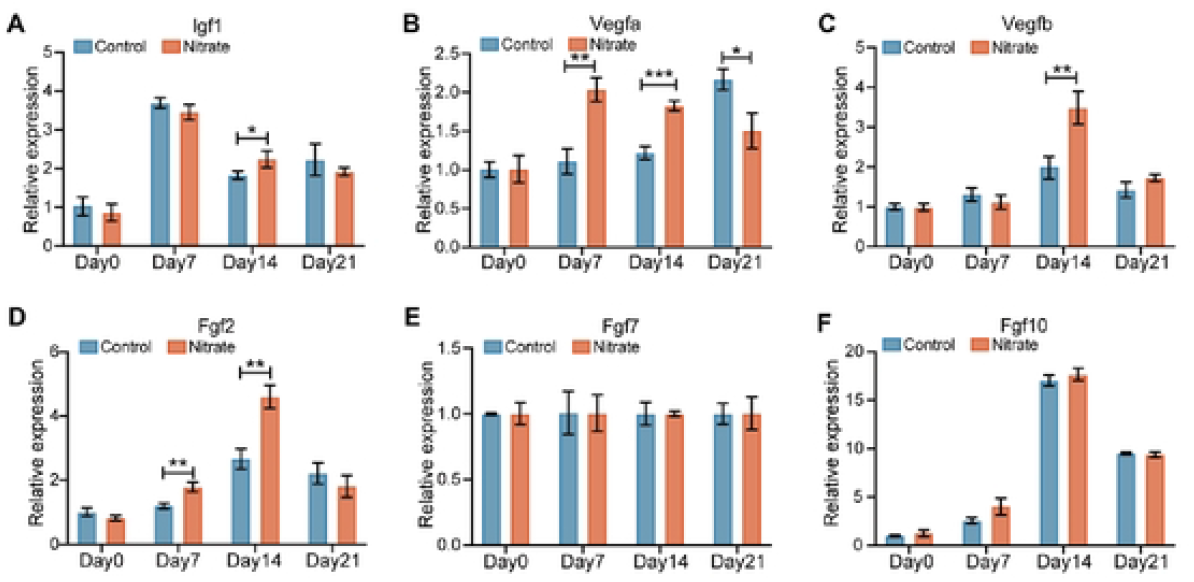
Nitrate increased the expression of Vegfa, Vegfb, Igf1 and Fgf2 in mice in promoting hair growth. (A-F) Relative mRNA expression of growth factors related to hair growth in mice treated with nitrate (10mM) or vehicle at days 0, 7, 14, and 21, (A) Igf1 (B) Vegfa; (C) Vegfb; (D) Fgf2; (E) Fgf7; (F) Fgf10;. All data are presented as the mean ± SD (n=4). * p < 0.05, ** p < 0.001, and *** p < 0.0001.

## Discussion

In this study, we explored the effect of nitrate administration on hair growth in vivo. Our results showed that nitrate supplementation had a positive effect on promoting hair growth in vivo. The active phase of the hair follicle cycle is accompanied by an increase in the number of follicles, resulting in an increase in the thickness of the subcutis layer between the dermis and panniculus carnosus. Immunohistochemistry, immunofluorescence, RT-PCR and western blot were used to detect the factors that influent by nitrate in hair growth in mice. We found hair promoting of nitrate was attributed to the enhancement of Wnt/β-catenin signal pathway, downregulation of TGF-β signal pathway and increased expression of significant growth factor, such as Vegf, Igf, and Fgf2 (Figure 7).

**Figure 7.**
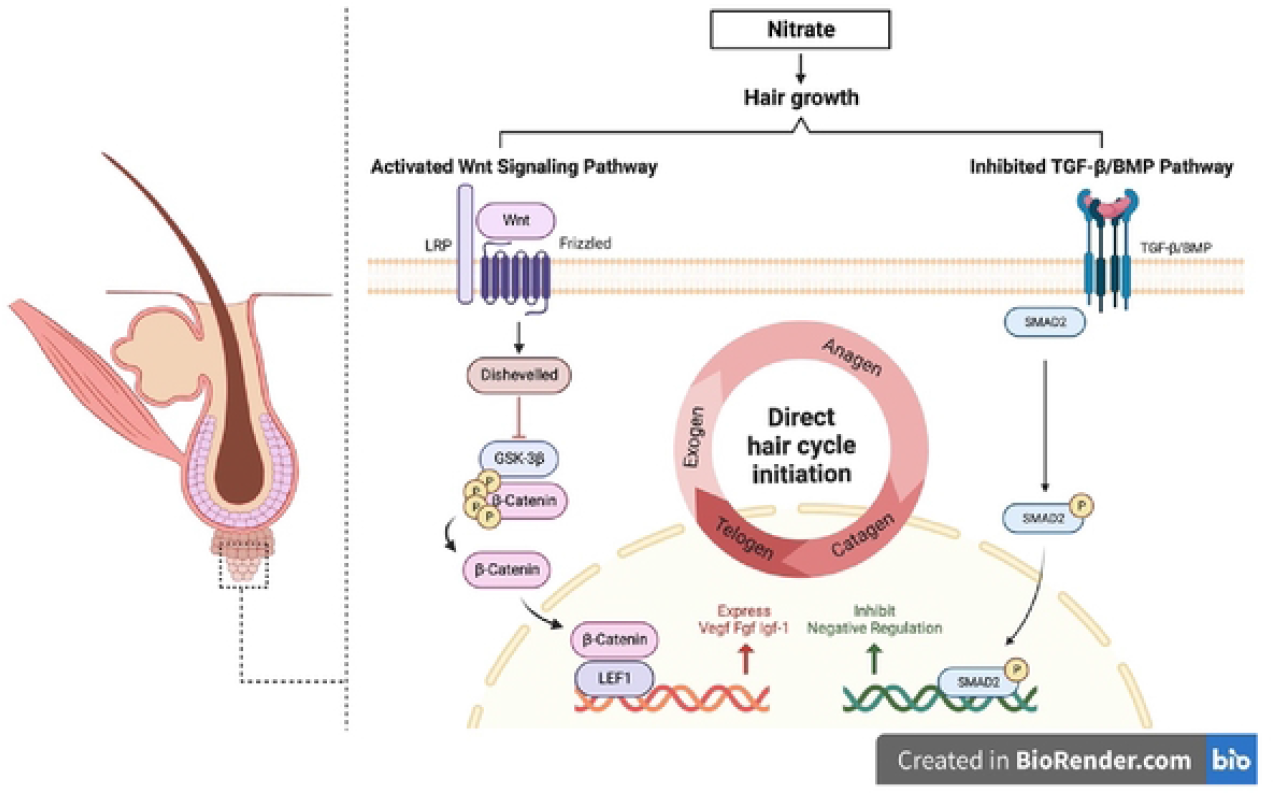
A proposed model for the role of nitrate in hair growth by activating Wnt/β-catenin and inhibiting TGF-β signaling pathway. In C57BL/6 mouse models, nitrate supplementation increased the level of β-catenin, the expression of its downstream molecules, as well as the expression of critical genes related to Wnt signaling and promote hair growth. Simultaneously, nitrate supplementation also decreased the level of TGF-β1 and the expression of Smad2, Bmp2 and Bmp6, and promote hair growth.

Now days, supplements or herbal remedies are becoming popular choices to prevent hair loss and promote hair growth. Nitrate, widely distributed in nature, and has been regarded as a potential carcinogen for decades[32]. Recently, researchers suggested that dietary nitrate shows protective effects in several physiological conditions [33]. Dietary nitrate supplementation always results in bioactive NO production through the nitrate-nitrite-NO pathway, which physiologically recycled in blood and tissues[12]. Moreover, NO was found playing various positive roles in the physiology of the skin[34, 35]. As a donor of NO, nitrate supplement (10mM) could promote hair growth in the present study. Actually, we designed 2mM (low) and 10mM (high) this two doses of nitrate to treat hair removed mice at first. But no significant differences were observed on hair growth in 2mM nitrate-treated mice (data not show). Nadeeka Bandara, et al., found that NO was a regulator of canonical Wnt/β-catenin signaling, and distinct concentration of exogenous NO differently modulates Wnt signaling[36]. Thus, we speculated that 10mM was an optimum concentration of exogenous NO to enhance Wnt/β-catenin signaling pathway and further promote hair growth. The biological effects of exogenous NO produced from nitrate supplementation toward hair growth must be studied deeply to obtain a better understanding of its role in the hair follicles and its microenvironments.

There are 2 forms of nitrate, organic and inorganic. Dietary nitrate is in the inorganic form. Particularly, inorganic nitrate lacks immunogenicity, and there are no subject to tolerance, no toxic, and no significant side effects at the doses employed[37]. Therefore, Dietary nitrate appears to comprise a safe and effective treatment for hair promoting. Previous studies on hair loss treatment mainly focused on topical treatments rather than oral supplementation. Patients often turn to oral supplements to address hair concerns as they are easily accessible, leading nitrate supplementation to become a good choice. It’s clear that after ingestion of a nitrate-rich diet, nitrate was absorbed and entered into the nitrate-enterosalivary circulation and nitrate-nitrite-NO pathway[38]. It has been reported that increased salivary nitrite production resulting from nitrate intake enhances oral NO production[39]. Therefore, it cannot be ignored that the beneficial effects of dietary nitrate are dependent on the processes in the oral cavity. Herein, oral administration of nitrate promoted hair growth might through this reabsorption step and induced oral NO, which is an important gaseous signaling molecule.

We investigated the effects of dietary nitrate on hair growth and regulation of classical pathways in mice. Wnt/β-catenin signaling has important roles during follicular growth and is related to cell proliferation of dermal papilla cells (DPCs)[40]. The results of the current study suggested that nitrate increased both the level of β-catenin and wnt3a in mice, and it is likely that the effects on Wnt/β-catenin signaling activation contributed to the increase in the number of hair follicular. Similarly, hedgehog (Shh) signaling also mediates β-catenin activity to direct hair follicle morphogenesis and hair cycle initiation. We also found the mRNA expression of shh was increased in nitrate group. DPCs secret diverse growth factors and other biomolecules, which are crucial in hair cycle. Significant genes, such as Fgf2, Vegf and Igf-1, involved in promoting growth or generation of hair, thus affecting follicular epithelium. In contrast, TGF-β1, related with catagen regulation, has been proposed to inhibit keratinocyte proliferation and induce apoptosis of hair follicle[41]. In this study, we observed nitrate upregulated he expression of Fgf2, Igf1 and Vegf, while downregulated the expression of TGF-β1. It is likely that Fgf2, Igf1, Vegf and Tgf-β1 may cross-talk to each other, as we know that the auto-induction and cross-induction mechanisms of growth factors are very effective in the growth of hair.

Overall, the present study demonstrated that nitrate supplementation stimulates hair growth by activating Wnt/β-catenin and inhibiting TGF-β signaling in vivo. Therefore, nitrate supplementation administration may be an effective approach for hair regrowth and additional studies are required to investigate the detail molecular mechanisms underlying these effects.

## Acknowledgments

This work was supported by funds from the Clinical Major Specialty Projects of Beijing (2-1-2-038 to LWB), and the Talent Introduction Project from Beijing Tiantan Hospital (RCYJ-2020-2025-LWB to LWB).

## Author contributions

L.W.B. and W.S.L. conceived the project, supervised the research, and revised the paper. Y.J.L. and L.S.L. performed most of the experiments, analyzed the data, and wrote the manuscript. W.J.C., D.L. and H.M.Q. assisted with the mouse experiments and data processing. Z.C.M. and C.F. participated in results and paper discussion. All authors have read and approved the final manuscript.

## Declare interests

The authors declare no conflict of interest.

---

**Figure S1. Evaluation of the biosafety of the nitrate.**

(A) Histopathological examination of liver and kidneys in two groups of mice treated with nitrate or vehicle. Scale bar = 100 μm.

(B) Measurements of body weight in the two groups of mice once a week after treatment. All data are presented as the mean ± SD (n=4). * p < 0.05, ** p < 0.001, and *** p < 0.0001.

**Figure S2. Nitrate increase the mRNA expression of β-catenin and its downstream genes.**

(A-D) Relative mRNA expression of genes related to β-catenin signaling in mice treated with nitrate (10mM) or vehicle at days 0, 7, 14, and 21; (A) β-catenin; (B) Gsk3β; (C) Lef1; (D) Ki67;

All data are presented as the mean ± SD (n=4). * p < 0.05, ** p < 0.001, and *** p < 0.0001.

**Figure S3. Nitrate increase the mRNA expression of critical genes related to Wnt signaling.**

(A-D) Relative mRNA expression of genes related to Wnt signaling in mice treated with nitrate (10mM) or vehicle at days 0, 7, 14, and 21; (A) Wnt4a; (B) Wnt5a; (C) Wnt5b; (D) Wnt7b.

All data are presented as the mean ± SD (n=4). * p < 0.05, ** p < 0.001, and *** p < 0.0001.

## Reference

1. Adil, A. and M. Godwin, The effectiveness of treatments for androgenetic alopecia: A systematic review and meta-analysis. J Am Acad Dermatol, 2017. 77(1):p. 136-141.e5.

2. Park, A.M., S. Khan, and J. Rawnsley, Hair Biology: Growth and Pigmentation. Facial Plast Surg Clin North Am, 2018. 26(4):p. 415–424.

3. Oh, J.W., et al., A Guide to Studying Human Hair Follicle Cycling In Vivo. J Invest Dermatol, 2016. 136(1):p. 34–44.

4. Stenn, K.S. and R. Paus, Controls of hair follicle cycling. Physiol Rev, 2001. 81(1):p. 449–494.

5. Paus, R. and K. Foitzik, In search of the “hair cycle clock”: a guided tour. Differentiation, 2004. 72(9-10): p. 489-511.

6. Phillips, T.G., W.P. Slomiany, and R. Allison, Hair Loss: Common Causes and Treatment. Am Fam Physician, 2017. 96(6):p. 371–378.

7. Strazzulla, L.C., et al., Alopecia areata: An appraisal of new treatment approaches and overview of current therapies. J Am Acad Dermatol, 2018. 78(1):p. 15–24.

8. Suchonwanit, P., S. Thammarucha, and K. Leerunyakul, Minoxidil and its use in hair disorders: a review. Drug Des Devel Ther, 2019. 13:p. 2777–2786.

9. Said, M.A. and A. Mehta, The Impact of 5α-Reductase Inhibitor Use for Male Pattern Hair Loss on Men’s Health. Curr Urol Rep, 2018. 19(8):p. 65.

10. Myint, P.K., et al., Fruit and vegetable consumption and self-reported functional health in men and women in the European Prospective Investigation into Cancer-Norfolk (EPIC-Norfolk): a population-based cross-sectional study. Public Health Nutr, 2007. 10(1):p. 34–41.

11. Hord, N.G., Y. Tang, and N.S. Bryan, Food sources of nitrates and nitrites: the physiologic context for potential health benefits. Am J Clin Nutr, 2009. 90(1):p. 1–10.

12. Lundberg, J.O., E. Weitzberg, and M.T. Gladwin, The nitrate-nitrite-nitric oxide pathway in physiology and therapeutics. Nat Rev Drug Discov, 2008. 7(2):p. 156–67.

13. Liu, H., et al., From nitrate to NO: potential effects of nitrate-reducing bacteria on systemic health and disease. Eur J Med Res, 2023. 28(1):p. 425.

14. Wolf, R., et al., Nitric oxide in the human hair follicle: constitutive and dihydrotestosterone-induced nitric oxide synthase expression and NO production in dermal papilla cells. J Mol Med (Berl), 2003. 81(2):p. 110–7.

15. Kim, H.J., et al., Non-thermal plasma promotes hair growth by improving the inter-follicular macroenvironment. RSC Adv, 2021. 11(45):p. 27880–27896.

16. Fujii, N., et al., Dietary nitrate supplementation increases nitrate and nitrite concentrations in human skin interstitial fluid. Nitric Oxide, 2023. 134-135: p. 10–16.

17. Huelsken, J., et al., beta-Catenin controls hair follicle morphogenesis and stem cell differentiation in the skin. Cell, 2001. 105(4):p. 533–45.

18. Ito, M., et al., Wnt-dependent de novo hair follicle regeneration in adult mouse skin after wounding. Nature, 2007. 447(7142):p. 316–20.

19. Houschyar, K.S., et al., Molecular Mechanisms of Hair Growth and Regeneration: Current Understanding and Novel Paradigms. Dermatology, 2020. 236(4):p. 271–280.

20. Kim, Y.E., et al., Costunolide promotes the proliferation of human hair follicle dermal papilla cells and induces hair growth in C57BL/6 mice. J Cosmet Dermatol, 2019. 18(1):p. 414–421.

21. Zhou, Y., et al., Identification of Hair Growth Promoting Components in the Kernels of Prunus mira Koehne and Their Mechanism of Action. Molecules, 2022. 27(16).

22. Kesika, P., et al., Role and Mechanisms of Phytochemicals in Hair Growth and Health. Pharmaceuticals (Basel), 2023. 16(2).

23. Zeng, Z., et al., Schizochytrium sp. Extracted Lipids Prevent Alopecia by Enhancing Antioxidation and Inhibiting Ferroptosis of Dermal Papilla Cells. Antioxidants (Basel), 2023. 12(7).

24. Rajan, P., et al., Effects of Cudrania tricuspidata and Sargassum fusiforme extracts on hair growth in C57BL/6 mice. Lab Anim Res, 2023. 39(1):p. 4.

25. Pérez-Mora, S., et al., BFNB Enhances Hair Growth in C57BL/6 Mice through the Induction of EGF and FGF7 Factors and the PI3K-AKT-β-Catenin Pathway. Int J Mol Sci, 2023. 24(15).

26. Li, S., et al., Liposomal honokiol promotes hair growth via activating Wnt3a/β-catenin signaling pathway and down regulating TGF-β1 in C57BL/6N mice. Biomed Pharmacother, 2021. 141:p. 111793.

27. Baek, Y.H., et al., Heat-Killed Enterococcus faecalis EF-2001 Induces Human Dermal Papilla Cell Proliferation and Hair Regrowth in C57BL/6 Mice. Int J Mol Sci, 2022. 23(10).

28. Jin, G.R., et al., Hair growth potential of Salvia plebeia extract and its associated mechanisms. Pharm Biol, 2020. 58(1):p. 400–409.

29. Junlatat, J. and B. Sripanidkulchai, Hair growth-promoting effect of Carthamus tinctorius floret extract. Phytother Res, 2014. 28(7):p. 1030–6.

30. Vañó-Galván, S., et al., Safety of low-dose oral minoxidil for hair loss: A multicenter study of 1404 patients. J Am Acad Dermatol, 2021. 84(6):p. 1644–1651.

31. Jung, H., et al., Mangifera Indica leaf extracts promote hair growth via activation of Wnt signaling pathway in human dermal papilla cells. Anim Cells Syst (Seoul), 2022. 26(3):p. 129–136.

32. Bryan, N.S., et al., Ingested nitrate and nitrite and stomach cancer risk: an updated review. Food Chem Toxicol, 2012. 50(10):p. 3646–65.

33. Ma, L., et al., Nitrate and Nitrite in Health and Disease. Aging Dis, 2018. 9(5):p. 938–945.

34. Heuer, K., et al., The topical use of non-thermal dielectric barrier discharge (DBD): nitric oxide related effects on human skin. Nitric Oxide, 2015. 44:p. 52–60.

35. Park, J., et al., Non-thermal atmospheric pressure plasma is an excellent tool to activate proliferation in various mesoderm-derived human adult stem cells. Free Radic Biol Med, 2019. 134:p. 374–384.

36. Bandara, N., et al., Molecular control of nitric oxide synthesis through eNOS and caveolin-1 interaction regulates osteogenic differentiation of adipose-derived stem cells by modulation of Wnt/β-catenin signaling. Stem Cell Res Ther, 2016. 7(1):p. 182.

37. Omar, S.A., E. Artime, and A.J. Webb, A comparison of organic and inorganic nitrates/nitrites. Nitric Oxide, 2012. 26(4):p. 229–40.

38. Zhang, H. and L. Qin, Positive feedback loop between dietary nitrate intake and oral health. Nutr Res, 2023. 115:p. 1–12.

39. Duncan, C., et al., Chemical generation of nitric oxide in the mouth from the enterosalivary circulation of dietary nitrate. Nat Med, 1995. 1(6):p. 546–51.

40. Zhang, Y., et al., Activation of beta-catenin signaling programs embryonic epidermis to hair follicle fate. Development, 2008. 135(12):p. 2161–72.

41. Inui, S., et al., Androgen-inducible TGF-beta1 from balding dermal papilla cells inhibits epithelial cell growth: a clue to understand paradoxical effects of androgen on human hair growth. Faseb j, 2002. 16(14):p. 1967–9.

